# The endoplasmic reticulum membrane protein complex (EMC) negatively regulates intestinal homeostasis through the Hippo signaling pathway

**DOI:** 10.1101/2022.01.25.477727

**Authors:** Lin Shi, Hubing Ma, Hang Zhao, Meifang Ma, Jinjun Wang, Ruiyan Kong, Zhengran Li, Rui Ma, Jian-Hua Wang, Shian Wu, Meng-qiu Dong, Zhouhua Li

## Abstract

Disruption of tissue homeostasis often results in many diseases. Balanced control of stem cell proliferation and differentiation underlines tissue homeostasis. However, how endogenous factors influence the proliferation and differentiation of intestinal stem cells (ISCs) under physiological conditions remains not fully understood. Here, we find that the endoplasmic reticulum membrane protein complex (EMC) negatively regulates ISC proliferation in adult *Drosophila* midgut. Compromising EMC function in progenitors leads to excessive ISC proliferation and intestinal homeostasis disruption. Mechanistically, the EMC complex associates with and stabilizes Hippo (Hpo), the key component of the Hpo signaling pathway. In the absence of the EMC complex, Yki (Yorkie) is activated to promote ISC proliferation. Furthermore, the role of the EMC complex in stem cell proliferation control is evolutionarily conserved. Thus, our study uncovers the molecular mechanism of the EMC complex in controlling stem cell proliferation. Our results provide new insight into the underlying mechanisms of how stem cell proliferation is properly controlled under physiological conditions.

## Introduction

Colorectal cancer is the fourth most deadly cancer in the world (after lung, liver, and stomach cancer). In 2020, there are about 915,880 new deaths from colorectal cancer globally, accounting for 9.2% of all cancer deaths (*Sung et al., 2021*). Previous studies have proved that intestinal epithelial cells have a high transformation rate, which compose a hot spot for the formation of malignant colorectal cancer (*Aran et al., 2016; Schwitalla et al., 2013; Zeuner et al., 2014*). Stem cells in adult tissues proliferate to produce differentiated cells, thereby maintaining tissue homeostasis. The homeostasis of the intestine is maintained by intestinal stem cells (ISCs). Therefore, in-depth study of the mechanism of ISC proliferation and differentiation is a key step to clarify the pathogenesis of colorectal cancer.

Studies have proved that the adult *Drosophila* intestine is a good model for studying tissue homeostasis and tumorigenesis (*Banerjee et al., 2019; Kim et al., 2020; Lucchetta and Ohlstein, 2012*). The *Drosophila* intestine is functionally equivalent to the small intestine in mammals. They are highly similar in development, cell type, and genetic regulation (*Casali and Batlle, 2009; Wang, 2010*). ISC in adult *Drosophila* intestine undergoes asymmetric division to produce a new stem cell and a daughter cell called enteroblast (EB) or enteroendocrine progenitor cell (EEP). Activation of Notch signaling in EB cells leads to further differentiation into enterocytes (ECs), while low level of Notch signaling causes EEP cells to divide once to produce two EE cells (*Chen et al., 2018; Fuss and Hoch, 2002; Noah and Shroyer, 2013; Tapanes-Castillo and Baylies, 2004*). In addition to the Notch signaling pathway, the proliferation and differentiation of ISC are also regulated by other evolutionarily conserved signaling pathways such as JAK/STAT, EGFR, Hippo (Hpo), Wnt, and BMP ((*Colombani and Andersen, 2020; Gervais and Bardin, 2017; Guo et al., 2016; Jiang et al., 2016; Joly and Rousset, 2020*) and references therein). Originally identified in *Drosophila* and evolutionarily conserved from fruit fly to mammals, the Hpo pathway plays an important regulatory role in cell proliferation and cell fate to control organ growth, regeneration, and tumorigenesis. In the canonical Hpo kinase cascade, the Hpo-Sav (Salvador) complex phosphorylates and activates the Wts (Warts)-Mats (mob as tumor suppressor) complex, activated Wts in turn phosphorylates and inactivates Yki (Yorkie) (*Chen et al., 2012; Misra and Irvine, 2018; Ren et al., 2010; Zheng and Pan, 2019*). Previous studies found that Hpo signaling negatively regulates ISC proliferation and intestinal homeostasis in *Drosophila (Karpowicz et al., 2010; Ren et al., 2010; Shaw et al., 2010; Staley and Irvine, 2010*). However, how Hpo signaling is regulated in progenitors remains not fully understood.

In eukaryotes, most protein synthesis and folding processes occur in the endoplasmic reticulum (ER). The evolutionarily conserved ER membrane protein complex (EMC) plays an important role in these processes (*Jonikas et al., 2009*). Analysis of the structure of human EMC complex showed that EMC contains 9 subunits (EMC1-8/9 and EMC10), in which EMC8 and EMC9 are ~40% identical (*Alvira et al., 2021; O’Donnell et al., 2020*). EMC has been proven to perform multiple functions in organisms: it mediates the insertion of the transmembrane domain (TMD) into the lipid bilayer and can act as a chaperone to participate in the folding of other types of membrane proteins in the ER (*Shurtleff et al., 2018; Tang et al., 2017; Volkmar and Christianson, 2020*). The lack of EMC will affect the physiological functions of organisms, such as ER stress, lipid homeostasis, tissue homeostasis, and autophagy (*Chitwood et al., 2018; Huang et al., 2021; Li et al., 2013; Umair et al., 2020; Volkmar et al., 2019; Yu et al., 2021; Zhu et al., 2020*). In addition, EMC10 governs male fertility via maintaining sperm ion balance in mice (*Zhou et al., 2018*), and EMC deficiency in *Drosophila* induces defective phototransduction and photoreceptor degeneration (*Satoh et al., 2015; Xiong et al., 2020*). However, whether and how EMC functions in ISC proliferation regulation remains unexplored.

In this study, we investigate the role of EMC in the proliferation of ISCs under physiological conditions in adult *Drosophila* intestine. We find that EMC is critical for ISC proliferation and midgut homeostasis. Our biochemical and genetic studies reveal that EMC interacts with and stabilizes Hpo to restrain excessive ISC proliferation, thereby maintaining intestinal homeostasis. Furthermore, the function of EMC in stem cell proliferation is evolutionarily conserved.

## Results

### EMC Complex Negatively Regulates ISC Proliferation

In order to identify endogenous factors that affect the homeostasis of adult *Drosophila* intestine, we conducted a large-scale RNAi screening using the *esgGal4, UAS-GFP, tubGal80^ts^* (*esg^ts^*) driver, which is expressed in progenitors (ISCs and EBs), in the posterior midgut (*Liu et al., 2021; Ma et al., 2016; Zhao et al., 2022*). The screen identified the ER membrane protein complex (EMC) as a candidate. Compared to control intestines in which *esg^+^* cells were infrequently observed in isolation or in clusters less than three cells, knocking down different subunits of EMC (EMC1, EMC3, EMC5, EMC6, and EMC7) led to dramatic increase in the number of *esg^+^* cells and intestinal homeostasis disruption (*Figure 1A–F, I*). The percentage of *esg^+^* cells (ratio of *esg^+^* cells to total cells) in these intestines was dramatically increased accordingly (*Figure 1J*). Consistently, quantitative RT-PCR (qRT-PCR) and western blotting (WB) results confirmed the knockdown efficiency (*Figure 1—figure supplement 1*). Meanwhile, significant increase in the number of mitotic cells was observed in *EMC1*-or *EMC3*-depleted intestines, suggesting that the dramatic increase in *esg>GFP* cells was caused by excessive ISC proliferation (*Figure 1K–N*). To further confirm that the defects are specific to EMC depletion, co-expression of an *UAS-EMC1-Flag* transgene with *EMC1^RNAi^* or an *UAS-EMC3-Myc* transgene with *EMC3^RNAi^* completely rescued the defects resulted from EMC1 or EMC3 depletion (*Figure 1G–J*). Finally, to confirm the RNAi results, we showed the size of *EMC1^GT1^* and *EMC3^e02662^* ISC MARCM clones was significantly increased compared to that of control clones (*Figure 1O–R*). Altogether, these data indicate that EMC negatively regulates ISC proliferation under physiological conditions.

**Figure 1.**
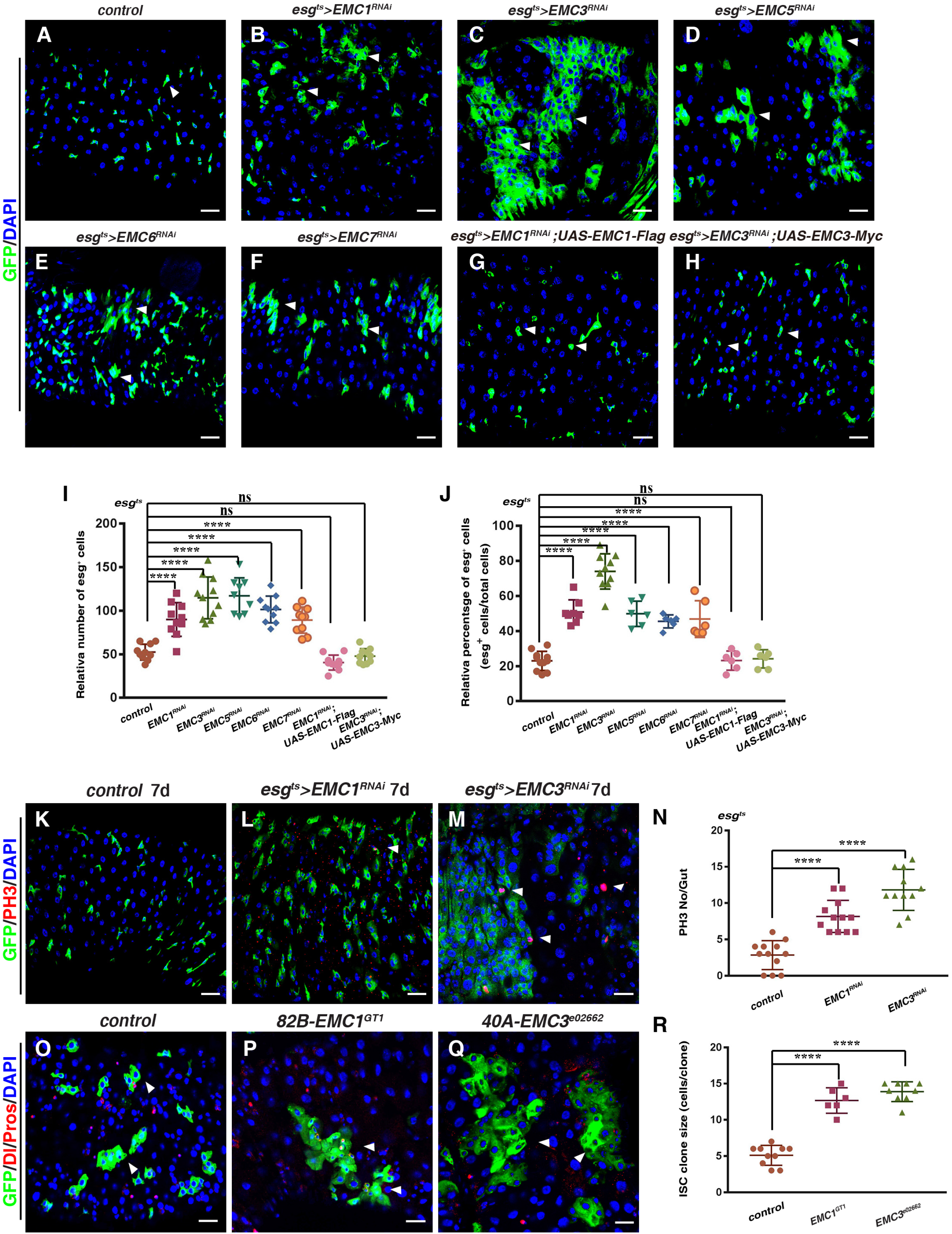
EMC Negatively Regulates ISC Proliferation. (A-H) Midguts of the following genotypes co-stained with DAPI (blue) driven by *esg^ts^* at 29°C for 7 days: WT (A), *UAS-EMC1^RNAi^* (B), *UAS-EMC3^RNAi^* (C), *UAS-EMC5^RNAi^* (D), *UAS-EMC6^RNAi^* (E), *UAS-EMC7^RNAi^* (F), *UAS-EMC1^RNAi^;UAS-EMC1-Flag* (G), and *UAS-EMC3^RNAi^;UAS-EMC3-Myc* (H). (I) Quantification *esg^+^* cell number per image (relative number). Mean ± SD is showed. n ≥ 9. ^ns^*p* > 0.05, \*\*\*\**p* < 0.0001. ns, not significant. (J) Quantification the percentage of *esg^+^* cell number per image. Mean ± SD is showed. n ≥ 6. ^ns^*p* > 0.05, \*\*\*\**p* < 0.0001. (K-M) Midguts of the following genotypes co-stained with DAPI (blue) and anti-PH3 (red) driven by *esg^ts^* at 29°C for 7 days: WT (K), *UAS-EMC1^RNAi^* (L), and *UAS-EMC3^RNAi^* (M). (N) Quantification of PH3^+^ cell number per midgut. Mean ± SD is showed. n ≥ 11. \*\*\*\**p* < 0.0001. (O-Q) MARCM clones co-stained with DAPI (blue) and anti-Dl/Pros (red). WT clone (O), *EMC1^GT1^* clone (P), and *EMC3^e02662^* clone (Q). (R) Quantification of the cell number per clone. Mean ± SD is showed. n ≥ 6. \*\*\*\**p* < 0.0001. Scale bars, 20 μm.

### Intestinal Homeostasis Is Disrupted in EMC-defective Intestines

Next, we characterized the identity of these extra *esg^+^* cells in *esg^ts^* > *EMC^RNAi^* intestines. We first stained the *esg^ts^* > *EMC^RNAi^* intestines with antibodies against Dl (an ISC marker) and Pros (an EE marker) after 7 days at 29°C. The number and percentage of Dl^+^ cells in *esg^ts^>EMC^RNAi^* intestines were significantly increased compared to those in control flies, whereas the number and the percentage of EEs were unaffected in these intestines (*Figure 2A–H*). The increase of Dl^+^ cells was independently verified with a *Dl-lacZ* reporter in *esg^ts^>EMC1^RNAi^* and *esg^ts^>EMC3^RNAi^* intestines (*Figure 2I–L*). Meanwhile, the number and percentage of EBs (by GBE+Su(H)-lacZ) were dramatically increased in *esg^ts^> EMC1^RNAi^* and *esg^ts^>EMC3^RNAi^* intestines compared to those in control flies (*Figure 2M–Q*). While none *esg^+^* cells in control flies were Pdm1^+^, many *esg^+^* cells within the *esg>GFP* clusters expressed Pdm1 in *esg^ts^>EMC^RNAi^* intestines (*Figure 2R–U*). Together, these data show that ISCs are over-proliferative in EMC-defective intestines, disrupting intestinal homeostasis.

**Figure 2.**
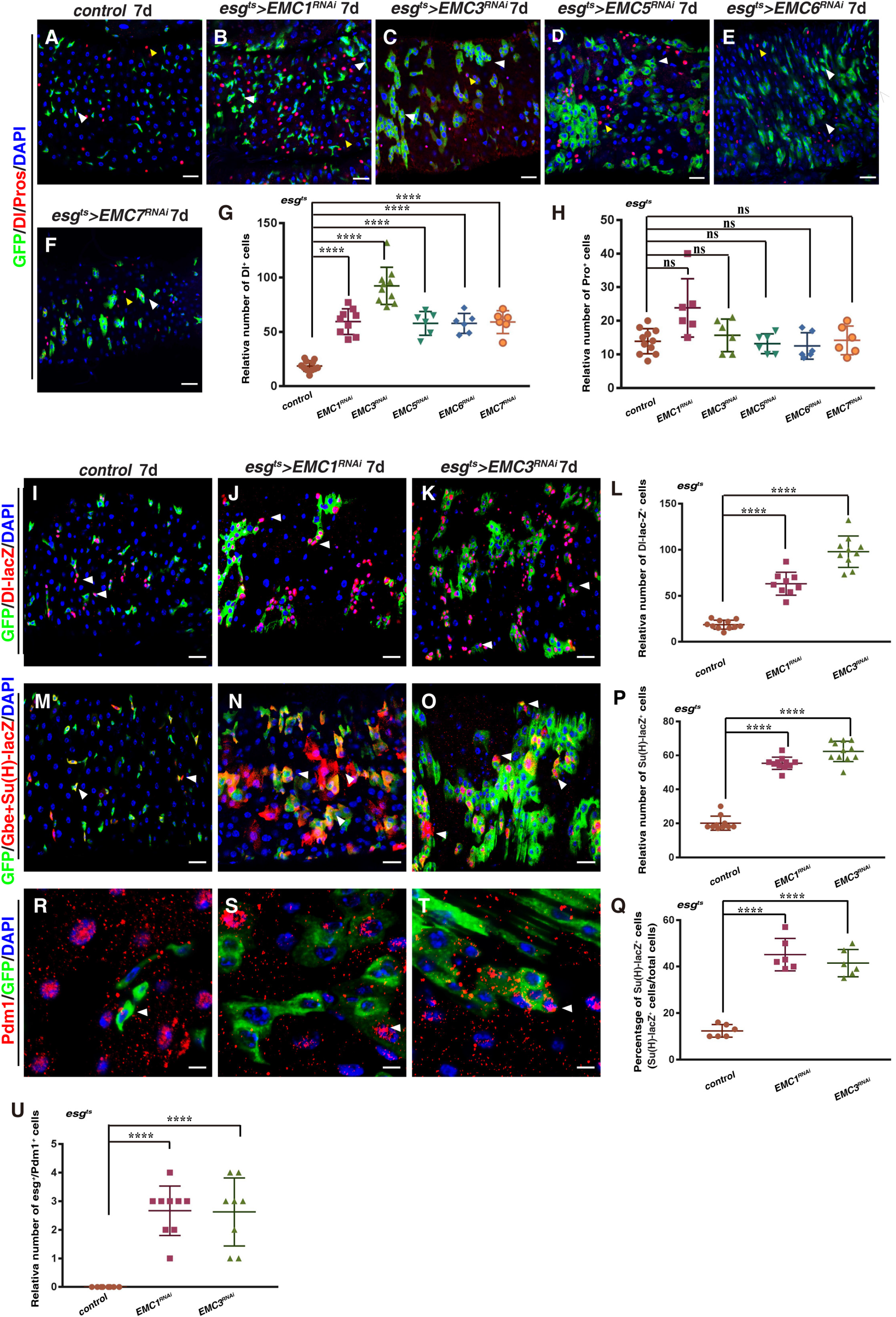
Intestinal Homeostasis Is Disrupted in *EMC-defective* Intestines. (A-H) Midguts of the following genotypes co-stained with DAPI (blue) and anti-Dl/Pros (red) driven by *esg^ts^* at 29°C for 7 days: WT (A), *UAS-EMC1^RNAi^* (B), *UAS-EMC3^RNAi^* (C), *UAS-EMC5^RNAi^* (D), *UAS-EMC6^RNAi^* (E), and *UAS-EMC7^RNAi^* (F). (G) Quantification of Dl^+^ cell number per image. Mean ± SD is showed. n ≥ 6. \*\*\*\**p*< 0.0001. (H) Quantification of Pros^+^ cell number per image. Mean ± SD is showed. n ≥ 6. ^ns^*p* > 0.05. (I-K) Midguts of the following genotypes stained with DAPI (blue) and anti-lacZ (red) driven by *esg^ts^, Dl-lacZ* at 29°C for 7 days: WT (I), *UAS-EMC1^RNAi^* (J), and *UAS-EMC3^RNAi^* (K). (L) Quantification of *Dl-lacZ*^+^ cell number per image. Mean ± SD is showed. n ≥ 9. \*\*\*\**p* < 0.0001. (M-O) Midguts of the following genotypes stained with DAPI (blue) and anti-lacZ (red) driven by *esg^ts^* at 29°C for 7 days: WT (M), *UAS-EMC1^RNAi^* (N), and *UAS-EMC3^RNAi^* (O). (P) Quantification of GBE+Su(H)-lacZ^+^ cell number per image. Mean ± SD is showed. n≥10. \*\*\*\**p* < 0.0001. (Q) Quantification the percentage of GBE+Su(H)-lacZ^+^ cell number per image. Mean ± SD is showed. n ≥ 6. \*\*\*\**p* < 0.0001. (R-T) Midguts of the following genotypes stained with DAPI (blue) and anti-Pdm1 (red) driven by *esg^ts^* at 29°C for 7 days: WT (R), *UAS-EMC1^RNAi^* (S), and *UAS-EMC3^RNAi^* (T). (U) Quantification of esg^+^/Pdm1^+^ cell number per image. Mean ± SD is showed. n ≥ 8. \*\*\*\**p* < 0.0001. Scale bars, 20 μm.

### Expression Pattern and Subcellular Localization of EMC

We then examined the expression pattern of EMC in the intestinal of *Drosophila* using the EMC1- and EMC3-specific polyclonal antibodies and showed that EMC1 and EMC3 were expressed in ISCs and other intestinal cell types (*Figure 2—figure supplement 1B, 2A–B*). Consistently with their ER localization, transiently expressed EMC1-GFP and EMC3-MYC localized to the ER in progenitors (*Figure 2—figure supplement 2C–D*). These results show that EMC1 and EMC3 localize in progenitors.

### EMC Associates with Hpo

To address how EMC regulates ISC proliferation, we performed coimmunoprecipitation (co-IP) and liquid chromatography-tandem mass spectrometry (LC-MS/MS) experiments to identify EMC1-interacting proteins by transiently expressing Flag-tagged EMC1. Most of the EMC subunits were identified, indicative of successful IP-MS (*Figure 3—figure supplement 3*). Among the candidate interactors, we identified Hpo, the core protein kinase in the Hpo signaling pathway. Previous studies have showed that the Hpo pathway is closely related to intestinal homeostasis regulation and tumorigenesis (*Chen et al., 2012; Harvey and Tapon, 2007; Jin et al., 2013; Karpowicz et al., 2010; Pan, 2010; Poernbacher et al., 2012; Ren et al., 2010; Shaw et al., 2010; Xu et al., 2019; Zeng and Hong, 2008*). To further confirm the association between EMC and Hpo, we performed co-IP experiments in *Drosophila* S2R^+^ cells. The results showed that EMC1 and Hpo reciprocally interact with each other in *Drosophila* S2R^+^ cells (*Figure 3A–B*). Moreover, *in vivo* co-IP results showed reciprocal interactions between transiently expressed EMC1 and transiently expressed Hpo (*Figure 3C–D*). In addition, transiently expressed EMC3 also interacts with transiently expressed Hpo *in vivo* (*Figure 3E*). Consistent with these data, both EMC1 and EMC3 co-localized with Hpo when they were transiently expressed in progenitors (*Figure 3—figure supplement 4A–B*). Furthermore, both endogenous EMC1 and EMC3 were co-expressed and co-localized with endogenous Hpo in the same intestinal cells (*Figure 3F–G*). Together, these results suggest an association between EMC and Hpo, indicating that EMC may regulate ISC proliferation through Hpo (and the Hpo signaling pathway).

**Figure 3.**
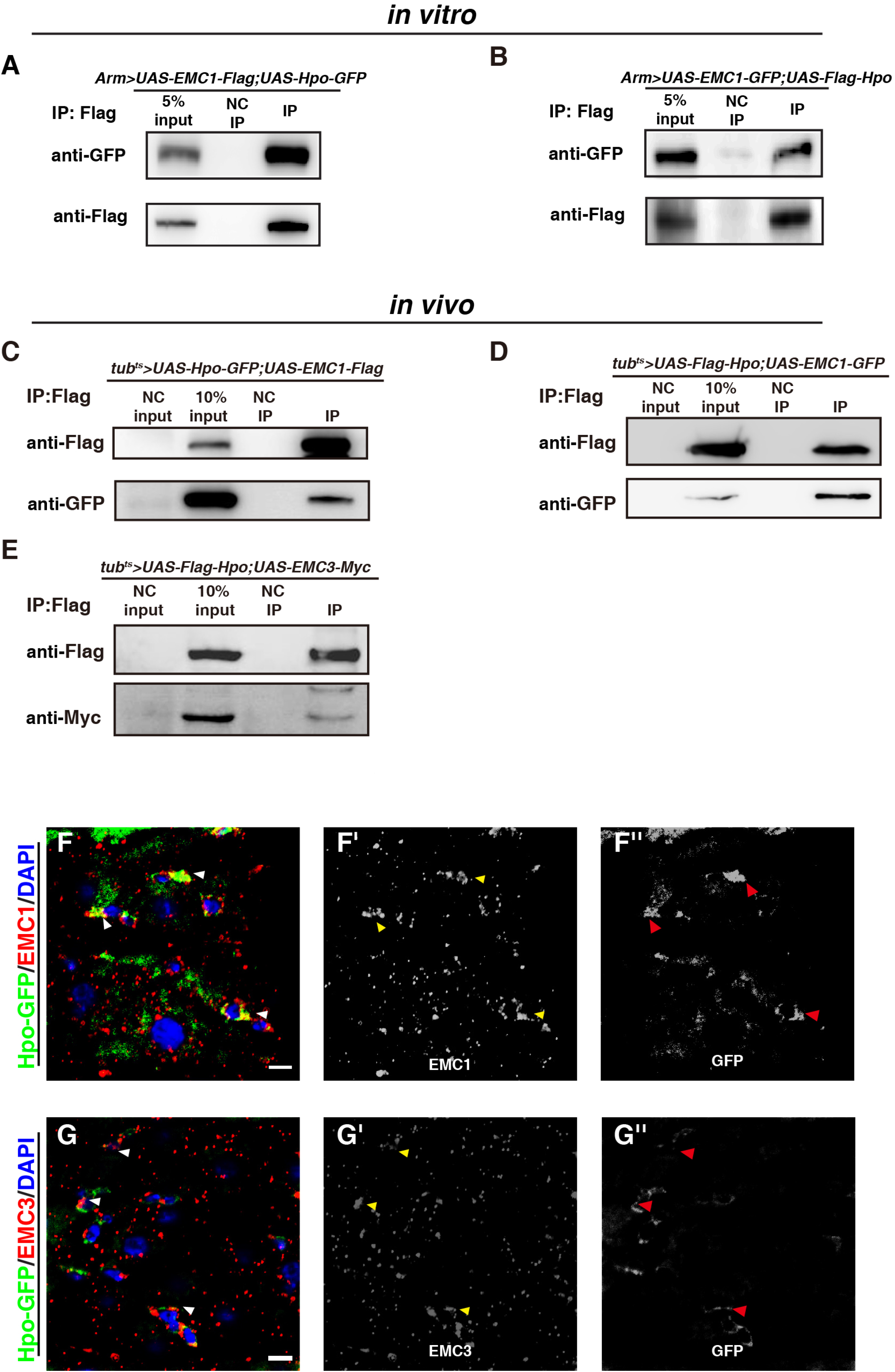
EMC Associates with Hpo. (A) co-immunoprecipitation (co-IP) results of *UAS-EMC1-Flag* and *UAS-Hpo-GFP* in S2R^+^ cells (Figure 3-source data 1, Figure 3-source data 2). NC: negative control. (B) co-IP results of *UAS-EMC1-GFP* and *UAS-Flag-Hpo* in S2R^+^ cells (Figure 3-source data 3, Figure 3-source data 4). (C) co-IP results of *UAS-EMC1-Flag* and *UAS-Hpo-GFP in vivo* (Figure 3-source data 5, Figure 3-source data 6). (D) co-IP results of *UAS-EMC1-GFP* and *UAS-Flag-Hpo in vivo* (Figure 3-source data 7, Figure 3-source data 8). (E) co-IP results of *UAS-Flag-Hpo* and *UAS-EMC3-Myc in vivo* (Figure 3-source data 9, Figure 3-source data 10). (F) Co-localization of EMC1 (red) and GFP-tagged Hpo (green) in intestines (arrowheads). EMC1 and Hpo-GFP channels are showed separately in black white. (G) Co-localization of EMC3 (red) and GFP-tagged Hpo (green) in progenitors (arrowheads). EMC3 and Hpo-GFP channels are showed separately in black white. Scale bars, 5 μm.

### Loss of EMC Leads to Inactivation of the Hpo Pathway

To further prove that EMC functions through the Hpo signaling pathway to regulate ISC proliferation, we examined the expression of several downstream targets of the Hpo signaling pathway. First, we found that the expression levels of *Diap1* (*Death associated inhibitor of apoptosis 1*), an anti-apoptotic gene, were significantly increased in *esg^ts^>EMC^RNAi^* intestines than those in control (*Figure 4A–D*) (*Huang et al., 2005; Yang and Choi, 2021*). Second, the expression levels of *bantam* microRNA (by *bantam-lacZ*) were significantly increased in *esg^ts^*>*EMC^RNAi^* intestines compared to those in control intestines (*Figure 4E–H*). Consistently, knocking down EMC led to a significant decrease in *bantam* sensor levels compared with control (*Figure 4—figure supplement 5A–D) (Brennecke et al., 2003*). Third, in the Hpo signaling pathway, a series of protein kinase cascade reactions lead to the final phosphorylation of Yki, phosphorylated Yki is inactivated as it cannot enter the nucleus to regulate the expression of downstream target genes (*Avruch et al., 2012; Huang et al., 2005*). The levels of phosphorylated Yki (p-Yki) were significantly reduced when EMC3 was systematically depleted, while the levels of total Yki proteins remained unchanged, indicative of activated Yki signaling in the absence of EMC (*Figure 4I*). Finally, we examined the activation of Yki signaling using a newly developed Yki-SPARK reporter (*Li et al., 2022*). Only a few Yki-SPARK droplets were detected in control midgut, in contrast, abundant Yki-SPARK droplets were observed in *esg^ts^>EMC^RNAi^* intestines (*Figure 4J-M*). Collectively, these observations indicate that Hpo signaling is inactivated and Yki is activated in the absence of EMC, which may be responsible for the defects resulted from EMC depletion.

**Figure 4.**
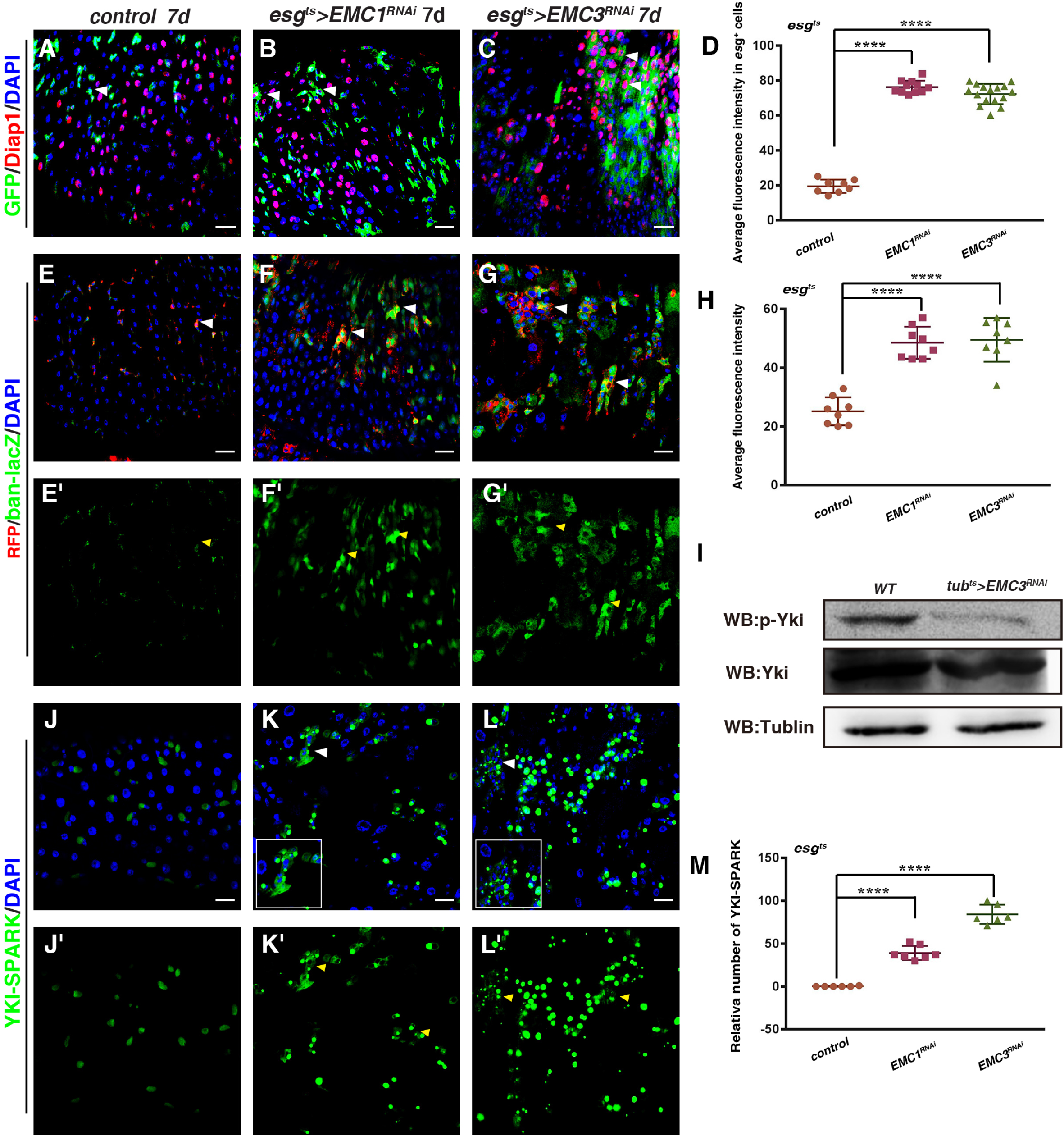
Loss of EMC Leads to Inactivation of the Hpo Pathway. (A-C) Midguts of the following genotypes stained with DAPI (blue) and anti-lacZ (red, by *Diap1-lacZ*) driven by *esg^ts^* at 29°C for 7 days: WT (A), *UAS-EMC1^RNAi^* (B), and *UAS-EMC3^RNAi^* (C). (D) Quantification average fluorescence intensity of *Diap1-lacZ* in intestines with indicated genotypes. Mean ± SD is showed. n ≥ 8. \*\*\*\**p* < 0.0001. (E-G) Midguts of the following genotypes stained with DAPI (blue) and anti-lacZ (green, by *ban-lacZ*) driven by *esg^ts^* at 29°C for 7 days: WT (E), *UAS-EMC1^RNAi^* (F), and *UAS-EMC3^RNAi^* (G). (H) Quantification average fluorescence intensity of *ban-lacZ* in intestines with indicated genotypes. Mean ± SD is showed. n ≥ 8. *\*\*\*\*p* < 0.0001. (I) The protein levels of p-Yki, Yki, and Tublin in WT and *tub^ts^ EMC3^RNAi^* flies at 29°C for 7 days (Figure-source data 1, Figure 4-source data 2, Figure 4-source data 3). (J-L) Yki-SPARK (green) in intestines with indicated genotypes. Yki-SPARK channel is showed separately. (M) Quantification of the Yki-SPARK number per image. Mean ± SD is showed. n ≥ 6. \*\*\*\**p* < 0.0001. Scale bars, 20 μm.

### EMC Stabilizes Hpo Protein

How does EMC affect the Hippo signaling pathway? We asked whether knocking down of EMC caused changes in Hpo protein levels. We examined the levels of endogenous Hpo protein by a reporter carrying a fosmid expressing GFP-labeled Hpo under its endogenous promoter (*Sarov et al., 2016*). Compared to the control, the levels of Hpo protein in progenitors were significantly reduced upon EMC knockdown (*Figure 5A–C*). Furthermore, the levels of Hpo protein were diminished upon systematic *EMC3* depletion, while the transcription levels of *hpo* were largely unaffected (*Figure 5D— figure supplement 5E*). These data indicate that EMC associates with and stabilizes Hpo protein to constrain ISC proliferation under physiological conditions. The 26S proteasome, one of the core elements of protein homeostasis regulation in eukaryotic cells, is responsible for the degradation of many proteins in various cellular pathways (*Bard et al., 2019; Bard et al., 2018; Coll-Martinez and Crosas, 2019*). We then examined the fate of Hpo protein in the absence of EMC. Interestingly, the levels of Hpo protein were significantly restored when the function of the 26S proteasome was further compromised (by depleting *Rpt5*) in *esg^ts^>EMC^RNAi^* intestines, indicating that Hpo protein is finally degraded by the 26S proteasome in the absence of EMC (*Figure 5C, E*). Altogether, these data indicate that EMC positively regulates the Hpo signaling pathway by stabilizing Hpo protein.

**Figure 5.**
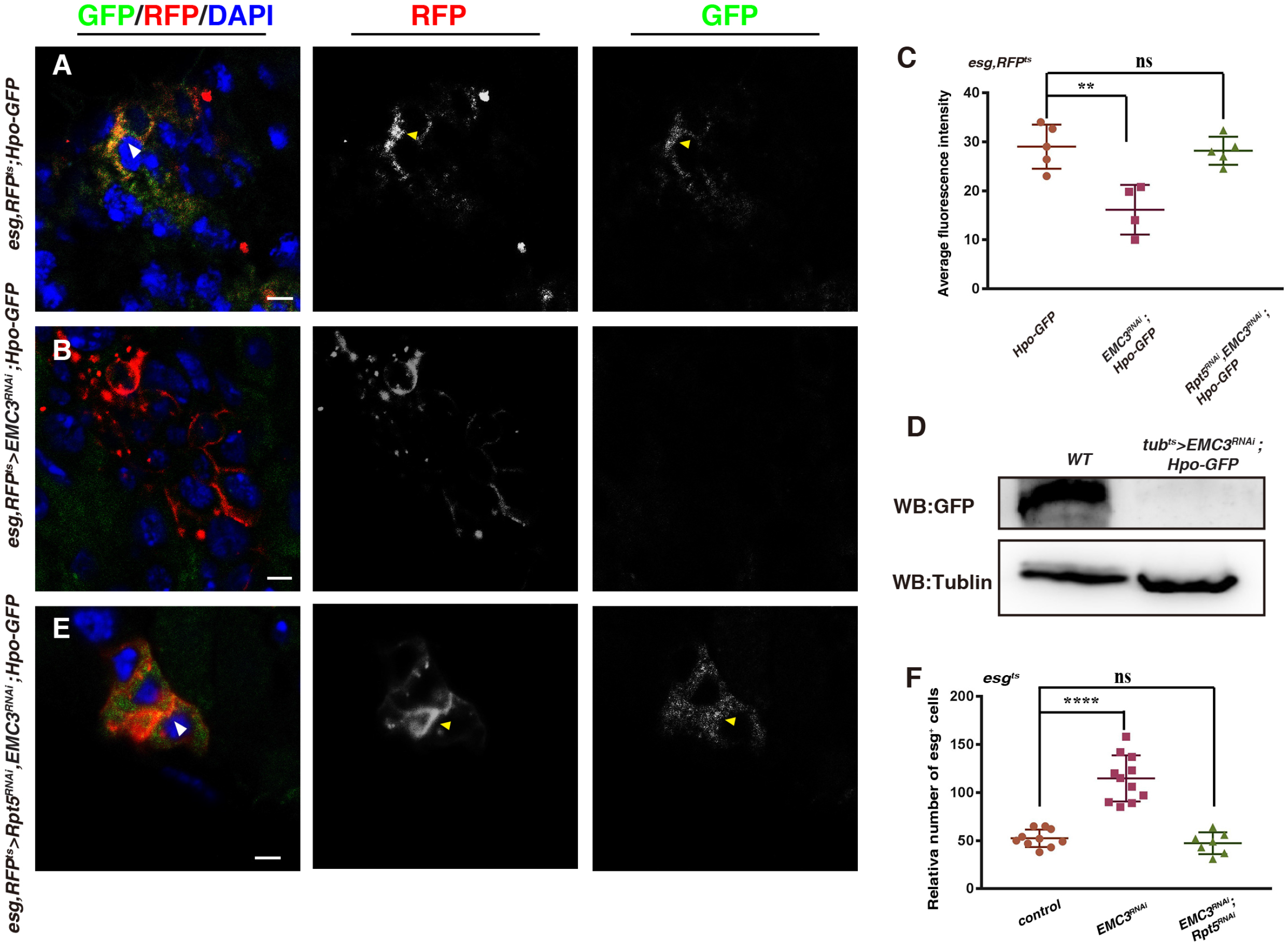
EMC Stabilizes Hpo Protein. (A-B) Midguts of the following genotypes stained with DAPI (blue): *esg-Gal4^ts^, UAS-RFP>w^RNAi^, Hpo-GFP* (A, control), *esg-Gal4^ts^, UAS-RFP>EMC3^RNAi^; Hpo-GFP* (B) at 29°C for 7 days (arrowheads). RFP and Hpo-GFP channels are showed separately in black white. (C) Quantification average fluorescence intensity of Hpo-GFP in intestines with indicated genotypes. Mean ± SD is showed. n ≥ 4. ^ns^*p* > 0.05, \*\**p* < 0.01. (D) The protein levels of Hpo-GFP in WT and *tub^ts^ EMC3^RNAi^* flies(Figure 5-source data 1, Figure 5-source data 2). (E) Hpo-GFP levels were significantly restored upon *Rpt5* depletion in *esg, RFP^ts^>EMC3^RNAi^* (F) Quantification the *esg^+^* cell number per image in intestines with indicated genotypes. Mean ± SD is showed. n ≥ 7. ^ns^*p* > 0.05, \*\*\*\**p* < 0.0001. Scale bars, 5 μm.

### Genetic Interactions between Hpo Signaling and EMC

Consistent with the notion that EMC functions through Hpo signaling to regulate ISC proliferation, *hpo* depletion or ectopic expression of constitutively active *yki* mimic those of EMC-defective intestines (*Figure 6—figure supplement 6A–F*). To functionally determine if defective Hpo signaling/ectopic Yki signaling is the cause for the defects associated with EMC depletion, we asked whether *esg^ts^>EMC^RNAi^* phenotypes could be suppressed by restoration of Hpo or inhibition of Yki. The increased number of *esg^+^* cells observed in *esg^ts^>EMC^RNAi^* intestines was effectively suppressed by co-depletion of Rpt5, which significantly restored the levels of Hpo protein (*Figure 5E–F*). Furthermore, co-expression of *Flag-Hpo* with *EMC1/3^RNAi^* completely suppressed the defects observed in *egs^ts^>EMC^RNAi^* intestines (*Figure 6A–F*). Similarly, co-expression of *yki^RNAi^* and *EMC3^RNAi^* also effectively inhibited the defects observed in *esg^ts^>EMC^RNAi^* intestines (*Figure 6G*). The number of GBE+Su(H)-lacZ^+^ and Dl^+^ cells in co-expressed intestines was restored to comparable levels as those in control flies (*Figure 6H—figure supplement 7A–D*). Consistently, the significant increase in mitotic ISCs was also completely suppressed in these intestines (*Figure 6—figure supplement 7E–H*). To further verify these results, we expressed *hpo* in *EMC1* or *EMC3* mutant cells and showed that ectopic expression of *hpo* effectively reduced the size of *EMC* mutant ISC MARCM clones (*Figure 6I–M, R*). In addition, knocking down of *yki* in *EMC1* or *EMC3* mutant clones significantly reduced the size of *EMC* mutant ISC MARCM clones (*Figure 6N–O,R Figure 6—figure supplement 7I–L,N*). Moreover, the size of *EMC3^RNAi^* MARCM clones was effectively suppressed by *yki* mutant (*Figure 6—figure supplement 7M–N*). These genetic interactions show that inactivation of Hpo signaling/activation of Yki is responsible for the defects resulted from *EMC* depletion. Altogether, our data demonstrate that EMC associates with and stabilizes Hpo protein in progenitors to ensure proper ISC proliferation, thereby maintaining midgut homeostasis.

**Figure 6.**
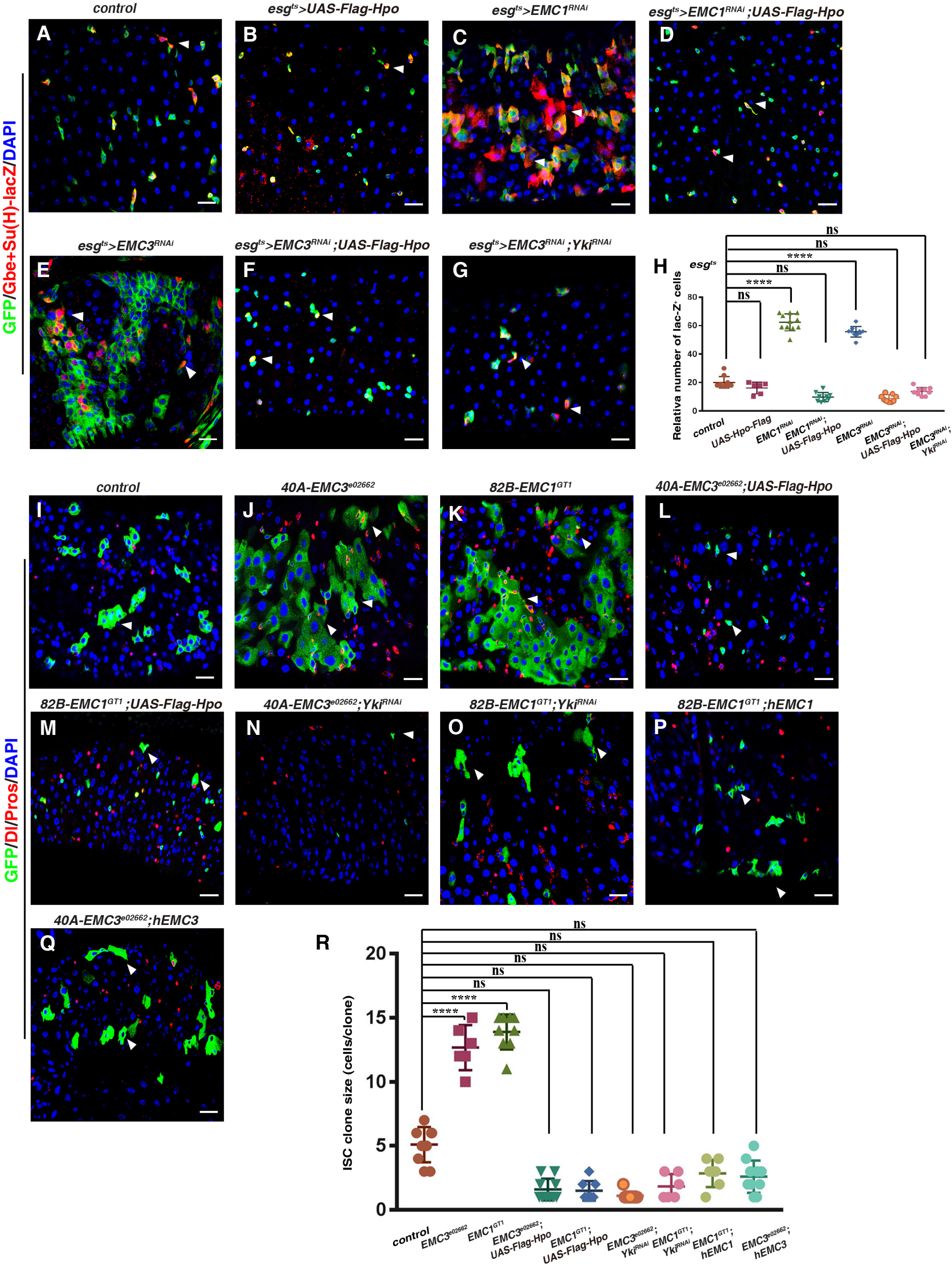
Genetic Interactions between Hpo Signaling and EMC. (A-G) Midguts of the following genotypes stained with DAPI (blue) and anti-lacZ (red, Su(H)-lacZ) driven by *esg^ts^* at 29°C for 7 days: WT (A), *UAS-Hpo-Flag* (B), *UAS-EMC1^RNAi^* (C), *UAS-EMC1^RNAi^;UAS-Hpo-Flag* (D), *UAS-EMC3^RNAi^* (E), *UAS-EMC3^RNAi^;UAS-Hpo-Flag* (F), and *UAS-EMC3^RNAi^;UAS-Yki^RNAi^* (G). (H) Quantification of *Gbe+Su(H)-lacZ^+^* cell number per image. Mean ± SD is showed. n ≥ 6. ^ns^*p* > 0.05, \*\*\*\**p* < 0.0001. (I-P) MARCM clones co-stained with DAPI (blue) and anti-Dl/Pros (red). WT clone (I), *EMC3^e02662^* clone (J), *EMC1^GT1^* clone (K), *EMC3^e02662^;UAS-Hpo-Flag* clone (L), *EMC1^GT1^;UAS-Hpo-Flag* clone (M), *EMC3^e02662^;UAS-Yki^RNAi^* clone (N), *EMC1^GT1^;UAS-Yki^RNAi^* clone (O), *EMC1^GT1^;UAS-hEMC1* clone (P), and *EMC3^e02662^;UAS-hEMC3* clone (Q). (R) Quantification of cell number per clone. Mean ± SD is showed. n ≥ 6. ^ns^*p* > 0.05, \*\*\*\**p* < 0.0001. Scale bars, 20 μm.

### The Function of EMC in ISC Proliferation Is Evolutionarily Conserved

Finally, we asked whether the function of EMC in ISC proliferation regulation was evolutionarily conserved. Ectopic expression of human *EMC1 (hEMC1)* effectively rescued the increased size of *Drosophila EMC1* mutant ISC MARCM clones (*Figure 6P, R*). Similarly, ectopic expression of human *EMC3 (hEMC3)* effectively rescued the increased size of *Drosophila EMC3* mutant ISC MARCM clones (*Figure 6Q–R*) These results show that the function of EMC in ISC proliferation regulation is evolutionarily conserved.

## Discussion

Activation of signaling pathways required for stem cell proliferation must be properly controlled to maintain tissue homeostasis. Here we demonstrate that EMC negatively regulates ISC proliferation. Our biochemical and genetic data show that EMC associates with and stabilizes Hpo protein to restrain ISC proliferation and maintain intestinal homeostasis under physiological conditions.

EMC plays important roles in protein synthesis and proper folding in the ER and it is related to a variety of physiological functions (*Bagchi et al., 2020; Chitwood and Hegde, 2019; Huang et al., 2021; Lahiri et al., 2014; Satoh et al., 2015*). However, previous studies have mostly focused on the role of EMC in the formation of the correct transmembrane structure (topology) of multi-transmembrane proteins or the function of EMC in lipid membrane transfer. In the absence of EMC, most of the signal anchors of the nascent chain cannot be inserted into correct topology, resulting in protein misfolding and degradation (*Kwon et al., 2019*). In *S. cerevisiae*, strains lacking multiple components of EMC reduce the transfer of phosphatidylserine (PS) from the ER to the mitochondria, leading to significantly reduced levels of PS and its derivative phosphatidylethanolamine (PE) in the mitochondria in these strains (*Lahiri et al., 2014*).

Different from its known functions in affecting the topological formation, folding, or transport of various membrane proteins (*Chitwood et al., 2018; O’Donnell et al., 2020; Volkmar and Christianson, 2020*), our results show that EMC also affects the stability of proteins destinated in cytoplasm such as Hpo, directly or indirectly. These data suggest that EMC may also affect various aspects of cytoplasmic protein processing in the ER, probably via facilitating the folding, processing, and/or transport of these proteins.

Previous studies indicated a close connection between EMC and ER-associated degradation (ERAD) components, which is related to ubiquitination and degradation of mis-folded proteins (*Christianson et al., 2012; Jonikas et al., 2009; Richard et al., 2013*). Consistent with previous report, we found that no ER stress responses (ER^UPR^) in progenitors were observed in the absence of EMC (*Wang et al., 2020*). These data indicate that in the absence of EMC, Hpo proteins are effectively processed by the ERAD pathway and the 26S proteasome, therefore, Hpo proteins are not accumulated in the ER and ER^UPR^ is not induced. Alternatively, as our recently study showed that no ER stress could be observed in progenitors when the key components of the ERAD pathway were compromised (*Liu et al., 2021*), the ER of progenitors can tolerate the temporary accumulation of Hpo proteins and can slowly remove these proteins from the ER without induction of ER^UPR^.

Intestinal stem cells are critical for intestinal homeostasis maintenance and imbalanced intestinal homeostasis is an important cause of intestinal tumors. Studies in recent years have showed that the Hpo signaling pathway plays an important role in ISC regulation, tissue regeneration, and cancer occurrence (*Harvey and Tapon, 2007; Huang et al., 2005; J1n et al., 2013; Karpowicz et al., 2010; Pan, 2010; Shaw et al., 2010; Zhang et al., 2009*). Our results here reveal the relationship between EMC deficiency and tumorigenesis in *Drosophila* adult intestines, indicating that EMC may be implicated in tumorigenesis by stabilizing Hpo protein to constrain stem cell proliferation and differentiation. Regarding the evolutionarily conserved function of EMC in stem cell regulation, and analyses of the ICGC (International Cancer Genome Consortium) database showed that aberrant expressions and/or mutations of *EMC* genes are observed in a variety of tumors, therefore, it will be interesting to investigate whether EMC deficiency is implicated in various human tumors in the future.

## Materials and methods

### Fly Lines and Cultures

Flies were maintained on standard media at 25°C. Crosses were raised at 18°C in humidity controlled incubators, or as otherwise noted. Flies hatched in 18°C incubators (2-3 days old) were picked and transferred to 29°C incubator, unless otherwise specified. Flies were transferred to new vials with fresh food every day, and dissected at time points specified in the text. In all experiments, only the female posterior midgut was analyzed. Information for alleles and transgenes used in this study can be found either in FlyBase, TRiP stock center at Tsinghua University (THU) or as noted: *esgGal4, UAS–CD8-GFP, tubGal80^ts^* (gift from N. Perrimon, *esg^ts^*), *esgGal4, UAS-CD8-RFP, tubGal80^ts^, EMC1^RNAi^* (HMS01055/THU1416 and VDRC9408), *EMC3^RNAi^* (HMS02242/THU4204), *EMC5^RNAi^* (VDRC48615), *EMC6^RNAi^* (VDRC49583), *EMC7^RNAi^* (VDRC8141), *EMC1^GT1^* (BL12852), *FRT40A-EMC3^e02662^* (Kyoto 114504), *FRT40A-EMC3/dPob^Δ4^* (gift from A. Satoh)(*Satoh et al., 2015*), *bantam* sensor (gift from D. Chen)(*Brennecke et al., 2003*), *bantam-lacZ* (gift from J. Shen), *yki^RNAi^* (HMS00041/THU0579), *FRT42D-yki^B5^* (gift from L. Zhang), *UAS-KDEL-GFP* (BL9898 and BL9899), *UAS-KDEL-RFP* (BL30910), *UAS-Flag-Hpo* (gift from L. Zhang) (*Jin et al., 2012*), *Hpo-GFP* (gift from J. Paster, VDRC318388)(*Sarov et al., 2016*), *Diap1-lacZ, UAS-Yki-SPARK* (gift from H. Huang)(*Li et al., 2022*), *Rpt5^RNAi^* (HMS00417/THU0868), *hpo^RNAi^* (THU551/HMS00006), *UAS-yki^CA^-V5* (BL28817), *w (white)^RNAi^* (BL33623, from TRiP at Harvard Medical School) was used as control.

### RNAi Knockdown and Overexpression Experiments

To address gene function in progenitors, *esgGal4, UAS-CD8-GFP, tubGal80^ts^* (*esg^ts^*) was used. The crosses (unless stated otherwise) were maintained at 18°C to bypass potential requirements during early developmental stages. 2-3 days old progeny with the desired genotypes were collected after eclosion and maintained at 29°C to inactivate Gal80^ts^ before dissection and immunostaining. The flies were transferred to new vials with fresh food every day. Both *UAS-dsRNA* and *UAS-shRNA* transgene stocks were used in this study. If possible, several dsRNA or shRNA lines were tested for each gene, and one or two RNAi lines were used for detailed study. The time points that the flies are analyzed/dissected were indicated in the text.

### Immunostainings and Fluorescence Microscopy

For standard immunostaining, intestines were dissected in 1 X PBS (10 mM NaH2PO4/Na2HPO4, 175 mM NaCl, pH7.4), and fixed in 4% paraformaldehyde for 25 min at room temperature. Samples were rinsed, washed with 1 X PBT (0.1% Triton X-100 in 1 X PBS) and blocked in 3% BSA in 1 X PBT for 45 min. Primary antibodies were added to the samples and incubated at 4°C overnight. The following primary antibodies were used: mouse mAb anti-Dl (C594.9B, 1:50, developed by S. Artavanis-Tsakonas, Developmental Studies Hybridoma Bank (DSHB)), mouse mAb anti-Prospero (MR1A, 1:100, developed by C.Q. Doe, DSHB), rabbit anti-β-glactosidase (lacZ, 1:5000, Cappel), mouse anti-β-glactosidase (lacZ, 1:1000, Cell Signaling), rabbit anti-PH3 (pSer10, 1:2000, Millipore, USA), rabbit anti-GFP (1:1000, Abcm), rabbit anti-Pdm1(1:200, gifts from B. Xiao and X. Yang)(*Dai et al., 2020; Yeo et al., 1995*), rabbit anti-EMC3(1:500), mouse anti-EMC3 (gift from A. Satoh) (*Satoh et al., 2015*), rabbit anti-EMC1(1:500), and mouse anti-Flag (1:1000, Sigma-Aldrich, USA). Secondary antibodies were incubated for 2 h at room temperature. DAPI (Sigma, 0.1 μg/mL) was added after secondary antibody staining. The samples were mounted in mounting medium (70% glycerol containing 2.5% DABCO). All images were captured by a Zeiss LSM780 inverted confocal microscope, and were processed in Adobe Photoshop and Illustrator.

### Constructs and Transgenes

The *EMC1* coding region was cloned into the EcoRI site of an *attB UAST-Flag* vector to generate *UAS-EMC1-FLAG.* The *EMC1* coding region was cloned into the EcoRI and XhoI sites of an *attB UAST-GFP* vector to generate *UAS-EMC1-GFP*. The *EMC3* coding region was cloned from LD37839 into the EcoRI and XbaI sites of an *attB UAST-MYC* vector to generate *UAS-EMC3-MYC*. The EMC3 coding region was cloned into the EcoRI and XhoI sites of an *attB UAST-GFP* vector to generate *UAS-EMC3-GFP*. The *hpo* coding region was cloned into the EcoRI and XhoI sites of an *attB UAST-GFP* vector to generate *UAS-Hpo-GFP*. The coding region of human *EMC1 (hEMC1)* was cloned into the XhoI and XbaI sites of an *attB UAST-V5* vector to generate *UAS-hEMC1-V5*. The coding region of human *EMC3* (*hEMC3*) was cloned into the EcoRI and XbaI sites of an *attB UAST-MYC* vector to generate *UAS-hEMC3-MYC*. Transgenic flies were obtained by standard P-element-mediated germline transformation carrying *attP* site at 86F or 36B.

### S2R+ Cell Transfection

S2R+ cells were cultured in 10 mL of heat-inactivated FBS medium supplemented with Penicillin-Streptomycin Solution at a ratio of 1:100. Transfection was carried out by PEI (polyethylenimine) transfection method when the cell density reached 2-4×10^6^ cells/cm. The medium was replaced with pre-warmed fresh complete medium 6-8 hours after transfection. The cells were harvested 48-72 hours after transfection.

### Generation of EMC1, EMC3, and Yki Antibodies

To generate polyclonal antibodies, two peptides of *Drosophila* EMC1 (EQKPRGDVKLLQVSGFADDSSDTAAA and SGSIVEMPWHLLDPRRPIASTTQ GREE) one peptide of *Drosophila* EMC3 (FNNEETGYFKTQKRAPVAQ), and two peptides of *Drosophila* Yki (cNPPSSHKPDDLEWYKIN and cMQTVHKKQRSYD VISPIQL) were synthesized. The peptides were immunized two rabbits respectively, and the antisera were affinity purified by BAM Biotech (BAMBIO, China).

### Co-Immunoprecipitation and Western Blotting

Fly tissues were lysed in RIPA buffer (50 mM Tris-HCl, pH8.0, 150 mM NaCl, 5 mM EDTA, pH8.0, 0.5% Triton X-100, 0.5% NP-40, 0.5% sodium deoxycholate, and complete protease inhibitor cocktail tablets (Roche)) on ice for 30 minutes. After centrifugation, lysates were then diluted ten-fold with RIPA buffer and subjected to immunoprecipitation using anti-FLAG M2 affinity gel (A2220; Sigma-Aldrich, USA). Immunocomplexes were collected by centrifugation and washed with 1 mL of RIPA buffer three times. Negative control IPs were performed for each immunoprecipitation experiment. For western blotting, immunoprecipitated proteins were separated in SDS-PAGE and then blotted onto PVDF membranes. Membranes were stained with primary antibody overnight at 4°C. Followed by washing, PVDF membranes were incubated with secondary antibodies conjugated with HRP, then the membranes were scanned using Luminescent Image Analyzer (GE, Sweden). Mouse anti-Flag (1:1000, Sigma-Aldrich, USA), mouse anti-Myc (9E10, 1:1000, Sigma-Aldrich, USA), rabbit anti-GFP (1:4000, Abcam, USA), rabbit anti-Yki S168 (p-Yki) (1:5000; gift from D. Pan)(*Dong et al., 2007*), rabbit anti-Yki (1:1000), rabbit anti-phospho-MST1 (Thr183)/MST2 (Thr180) (p-Hpo, 3681S, 1:1000, Cell Signaling, USA), rabbit anti-EMC3 (1:1000, this study), rabbit anti-EMC1(1:500, this study), and mouse monoclonal anti–αTubulin or anti-βTubulin (1:1000, Abbkine, USA) antibodies were used.

### MS Sample Preparation

Immunoprecipitated proteins from whole body of *tub^ts^>EMC1-Flag* flies were precipitated with 25% trichloroacetic acid (TCA) for at least 30 min on ice. Protein pellets were washed twice with 500 μL ice-cold acetone, air dried, and then resuspended in 8 M urea, 100 mM Tris, pH8.5. After reduction (5 mM TCEP, RT, 20 min) and alkylation (10 mM iodoacetamide, RT, 15 min in the dark), the samples were diluted to 2 M urea with 100 mM Tris, pH8.5 and digested with trypsin at 1/50 (w/w) enzyme/substrate ratio at 37°C for 16-18 hr. Digestion was then stopped by addition of formic acid to 5% (final concentration).

### LC-MS/MS Analysis

All samples were analyzed using an EASY-nLC 1000 system (Thermo Fisher Scientific, Waltham, MA) interfaced with a Q-Exactive mass spectrometer (Thermo Fisher Scientific). Peptides were loaded on a trap column (75 μm ID, 4 cm long, packed with ODS-AQ 12 nm-10 μm beads) and separated on an analytical column (75 μm ID, 12 cm long, packed with Luna C18 1.9 μm 100 Å resin) with a 60 min linear gradient at a flow rate of 200 nl/min as follows: 0-5% B in 2 min, 5-30% B in 43 min, 30-80% B in 5 min, 80% B for 10 min (A = 0.1% FA, B = 100% ACN, 0.1% FA). Spectra were acquired in data-dependent mode: the top ten most intense precursor ions from each full scan (resolution 70,000) were isolated for HCD MS2 (resolution 17,500; NCE 27) with a dynamic exclusion time of 30 s. The AGC targets for the MS1 and MS2 scans were 3e6 and 1e5, respectively, and the maximum injection times for MS1 and MS2 were both 60 ms. Precursors with 1+, more than 7+ or unassigned charge states were excluded.

### Database Search

MS data were searched against a Uniprot *Drosophila melanogaster* protein database (database ID number of UP000000803) using ProLuCID with the following parameters: precursor mass tolerance, 3 Da; fragment mass tolerance 20 ppm; peptide length, minimum 6 amino acids and maximum 100 amino acids; enzyme, Trypsin, with up to three missed cleavage sites (*Xu et al., 2015*). Results were filtered by DTASelect requiring FDR<1% at the peptide level and spectra count >= 2 (*Tabb et al., 2002*). Proteins identified from the negative control and Flag-Yun IP were contrasted by Contrast (Tabb et al., 2002).

### Data Analysis

PH3 numbers were scored manually under Zeiss Imager Z2/LSM780 microscope for indicated genotypes. To determine the number of indicated cells per confocal image (the relative number), confocal images of 40 X lens/1.0 zoom from a defined posterior midgut region of different genotypes indicated were acquired. The relative number of indicated cell types (such as progenitor cells and *Dl-lacZ^+^* cells), PH3^+^ cells, and the YKi-SPARK puncta per cell were determined using Image-Pro Plus software from each confocal image. The percentage of indicated cells was the ratio of the number of these cells per image to the total cell number in this image. Fluorescence intensity (IOD) was measured by ImageJ software for indicated genotypes. The number of intestines scored is indicated in the text. Statistical analysis was done by the unpaired Student’s *t*-test using GraphPad Prism 8 software. The graphs were further modified using Adobe Photoshop and Illustrator. \**p* < 0.05, \*\**p* < 0.01, \*\*\**p* < 0.001, \*\*\*\**p* < 0.0001.

### Real-time qPCR

Total RNA was extracted from 30 flies or guts using TRIzol (Invitrogen) according to manufacturer’s instructions. RNA was cleaned using RNAeasy (QIAGEN), and complementary DNA (cDNA) was synthesized using the iScript cDNA synthesis kit (Bio-Rad). Quantitative PCR was performed using the iScript one step RT-PCR SYBR green kit (Bio-Rad). RT-qPCR was performed in duplicate for each of three independent biological replicates. All results are presented as mean ± SD of the biological replicates. The ribosomal gene *RpL11* was used as the normalization control.

## ACKNOWLEDGEMENTS

We are grateful to Norbert Perrimon, Sarah Bray, Rongwen Xi, Jose Pastor, Xiaohang Yang, Xiaolin Bi, Lei Zhang, Hai Huang, Jie Shen, Dahua Chen, Xing Wang, Steve Cohen, Duojia Pan, and Yu Cai for generous gifts of reagents, the Bloomington Stock Center, VDRC, NIG-FLY Center, TRiP at Harvard Medical School (NIH/NIGMS R01-GM084947), and Tsinghua Fly Center (THFC) for fly stocks, DSHB for antibodies, and DGRC for cDNA clones. This work is supported by grants from the National Natural Science Foundation of China (Nos. 31972893, 92054109, and 31471384) and Beijing Municipal Commission of Education (No. KZ201910028040). We declare no conflicts of interest.

## Supplementary Information

### Supplementary Figures

**Figure S1.**
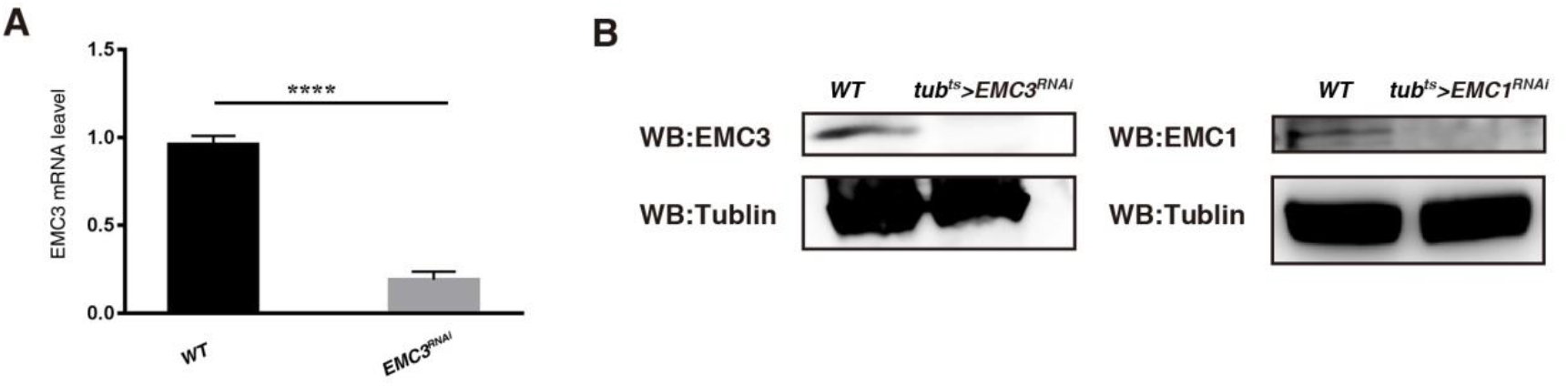
RNAi Knockdown Efficiency and Antibody Specificity. (A) The mRNA levels of *EMC3* in *WT* and *tub^ts^>EMC3^RNAi^* flies at 29°C for 7 days. Mean ± SD is showed. \*\*\*\**p* < 0.0001 (Figure1-figure supplement 1 -source data 1, Figure1-figure supplement 1 -source data 2, Figure1-figure supplement 1-source data 3, Figure1-figure supplement 1 -source data 4). (B) The protein levels of EMC3 and EMC1 are dramatically decreased in *tub^ts^>EMC3^RNAi^* and *tub^ts^>EMC1^RNAi^* flies.

**Figure S2.**
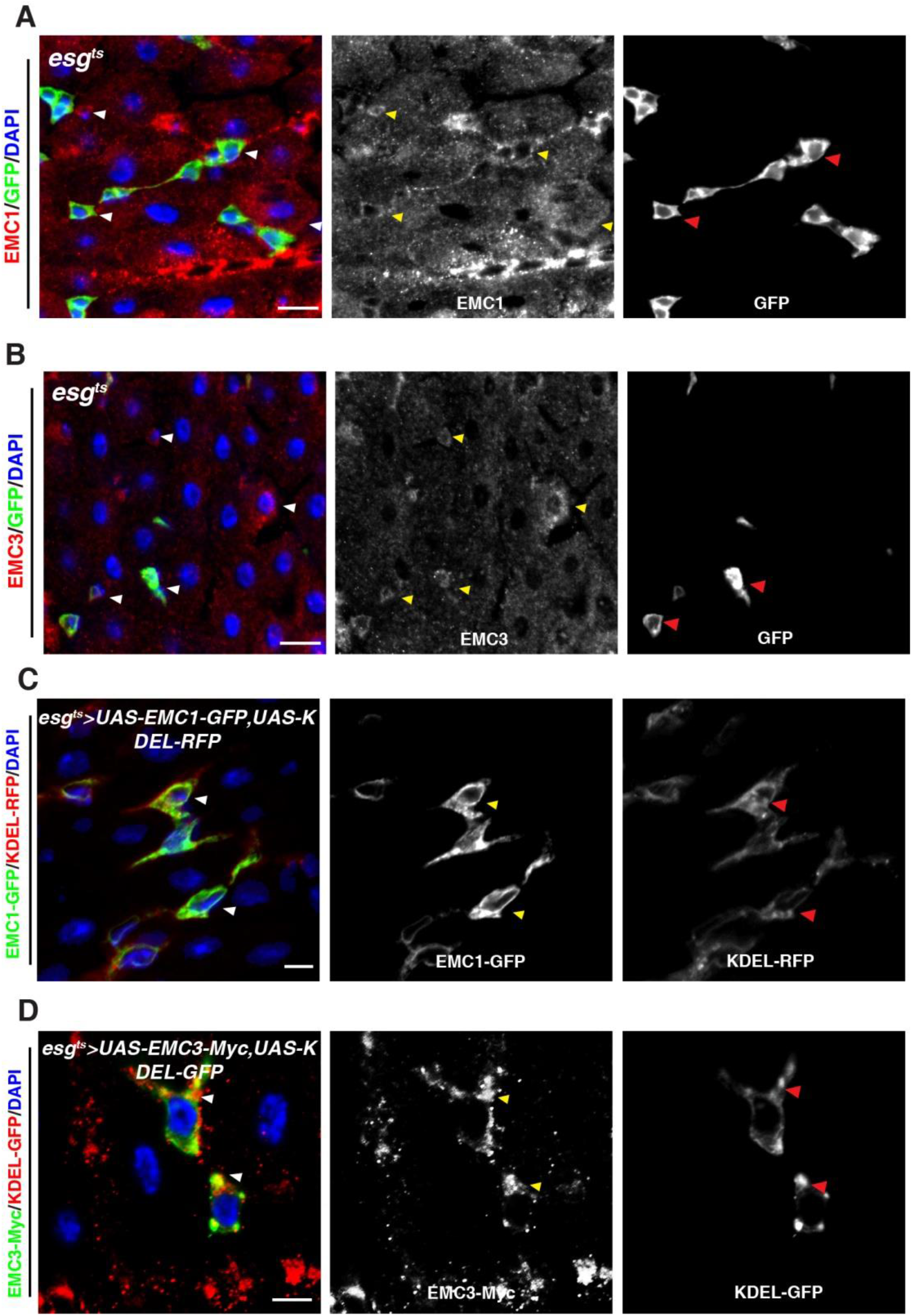
Expression Pattern and Subcellular Localization of EMC. (A) Midguts stained with anti-EMC1 antibody (red). EMC1 is expressed in progenitors (green) and other intestinal cell types (arrowheads). EMC1 and GFP channels are showed separately in black white. (B) Midguts stained with anti-EMC3 antibody (red). EMC3 is expressed in progenitors (green) and other intestinal cell types (arrowheads). EMC3 and GFP channels are showed separately in black white. (C) Co-localization of GFP-tagged EMC1 (green) and RFP-tagged KDEL (red, the ER) in progenitors (arrowheads). EMC1-GFP and KDEL-RFP channels are showed separately in black white. (D) Co-localization of Myc-tagged EMC3 (red) and GFP-tagged KDEL (green, the ER) in progenitors (arrowheads). EMC3-Myc and KDEL-GFP channels are showed separately in black white. Scale bars, 20μm (A, B), 10 μm (C), and 5 μm (D).

**Figure S3.**
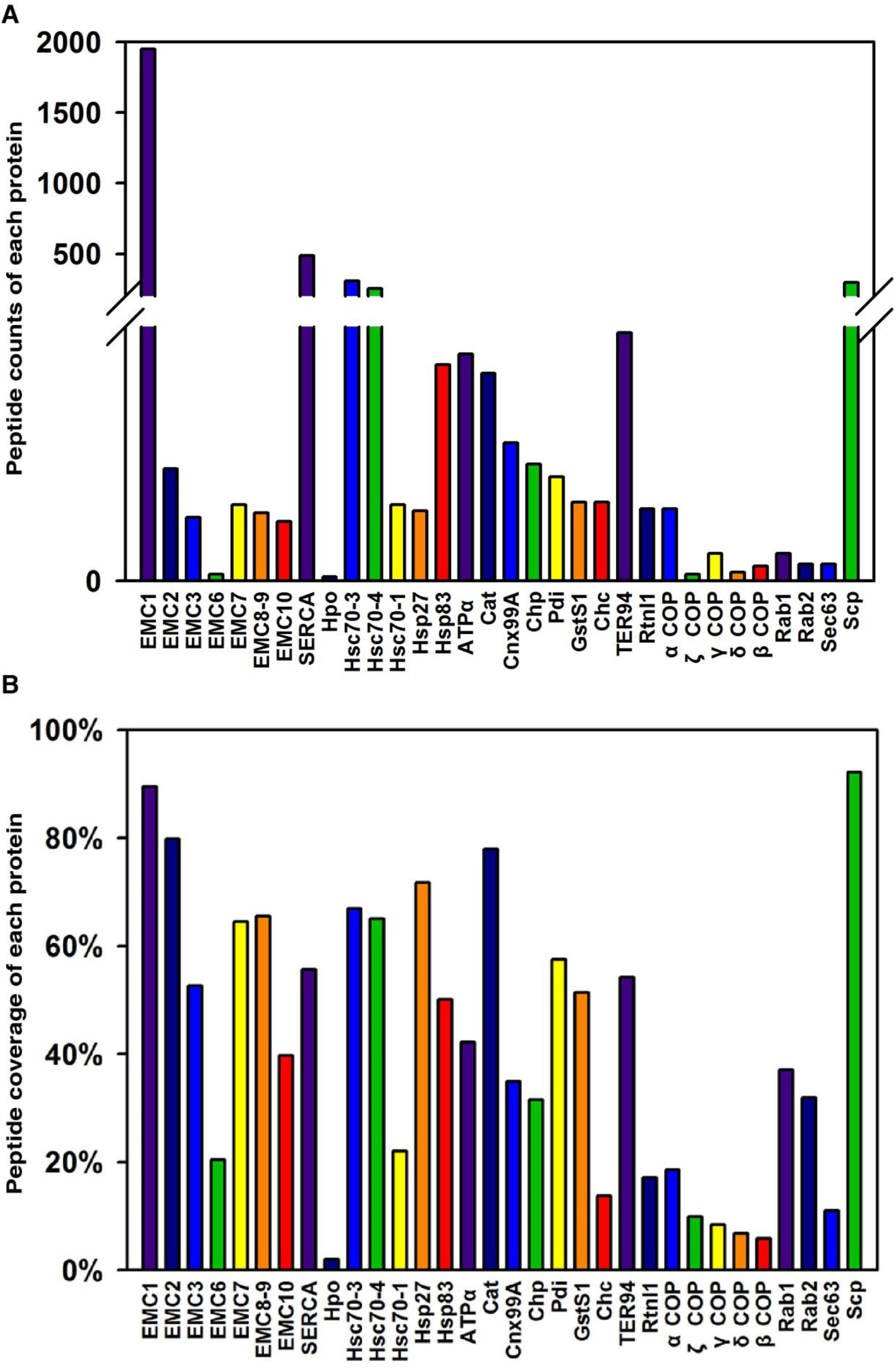
EMC-interacting Proteins Identified by Mass Spectrometry. (A) Peptide counts of selected proteins identified by mass spectrometry. (B) Peptide coverage of selected protein identified by mass spectrometry.

**Figure S4.**
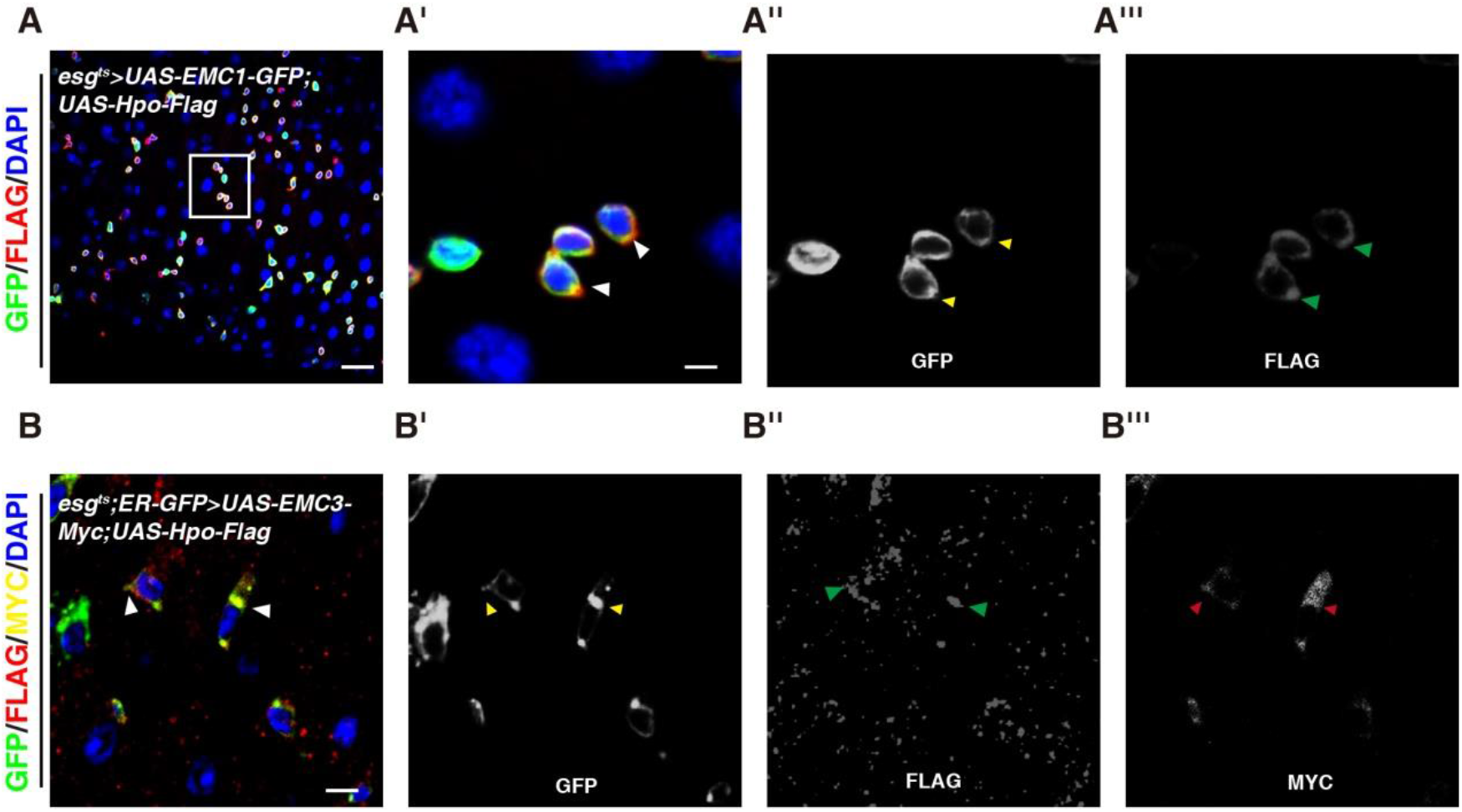
EMC Associates with Hpo. (A) Co-localization of GFP-tagged EMC1 (green) and FLAG-tagged Hpo (red) in progenitors (arrowheads). EMC1-GFP and Flag-Hpo channels are showed separately in black white. (B) Co-localization of ER-GFP (green), Myc-tagged EMC3 (yellow) and Fla-tagged Hpo (red) in progenitors (arrowheads). ER-GFP, EMC3-Myc, and Flag-Hpo channels are showed separately in black white. Scale bars, 20 μm (A); 5 μm (A’, A’’, A’’’, B, B’, B’’, B’’’).

**Figure S5.**
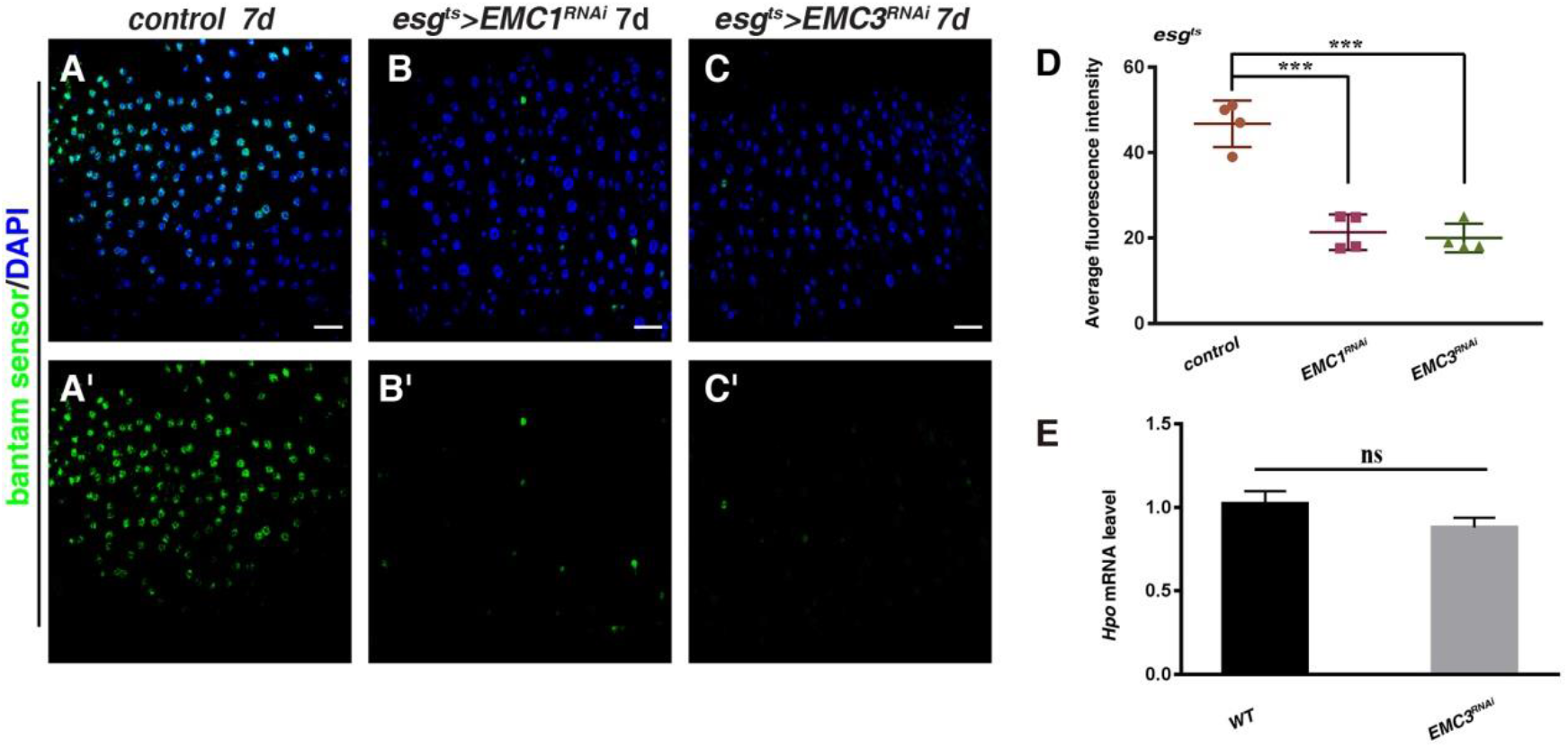
Loss of EMC Leads to Inactivation of the Hpo Pathway. (A-C) *bantam* sensor in the following intestines: *esg^ts^, ban sensor>w^RNAi^* (A, control), *esg^ts^, ban sensor>UAS-EMC1^RNAi^* (B), and *esg^ts^, ban sensor>UAS-EMC3^RNAi^* (C). (D) Quantification average fluorescence intensity of ban sensor in intestines with indicated genotypes. Mean ± SD is showed. n ≥ 4. \*\*\**p* < 0.001. (E) The mRNA level of *Hpo* is unaffected by EMC3 depletion. ^ns^*p* > 0.05. Scale bars, 20 μm.

**Figure S6.**
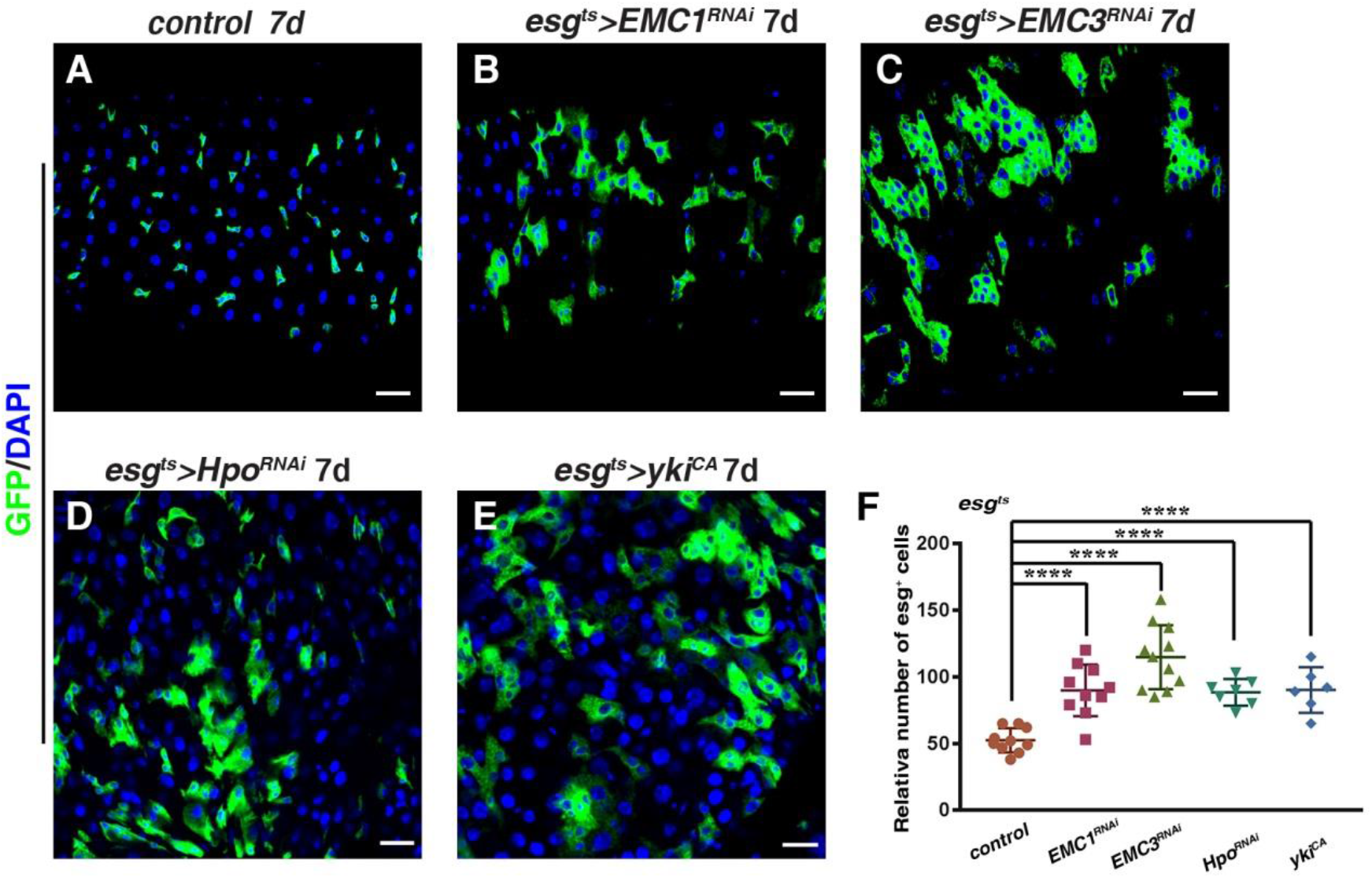
*hpo* Depletion and Constitutively Active *yki* Mimic the EMC-defective Intestines. (A-E) Midguts of the following genotypes stained with DAPI (blue): *esg^ts^>w^RNAi^* (A), *esg^ts^>EMC1^RNAi^* (B), *esg^ts^>EMC3^RNAi^* (C), *esg^ts^HP0^RNAi^* (D), *esg^ts^>yki^CA^* (E). (F) Quantification of the *esg^+^* cell number in intestines with indicated genotypes. Mean ± SD is showed. n ≥ 6. ^ns^*p* > 0.05, \*\*\*\**p* < 0.0001. Scale bars, 20 μm.

**Figure S7.**
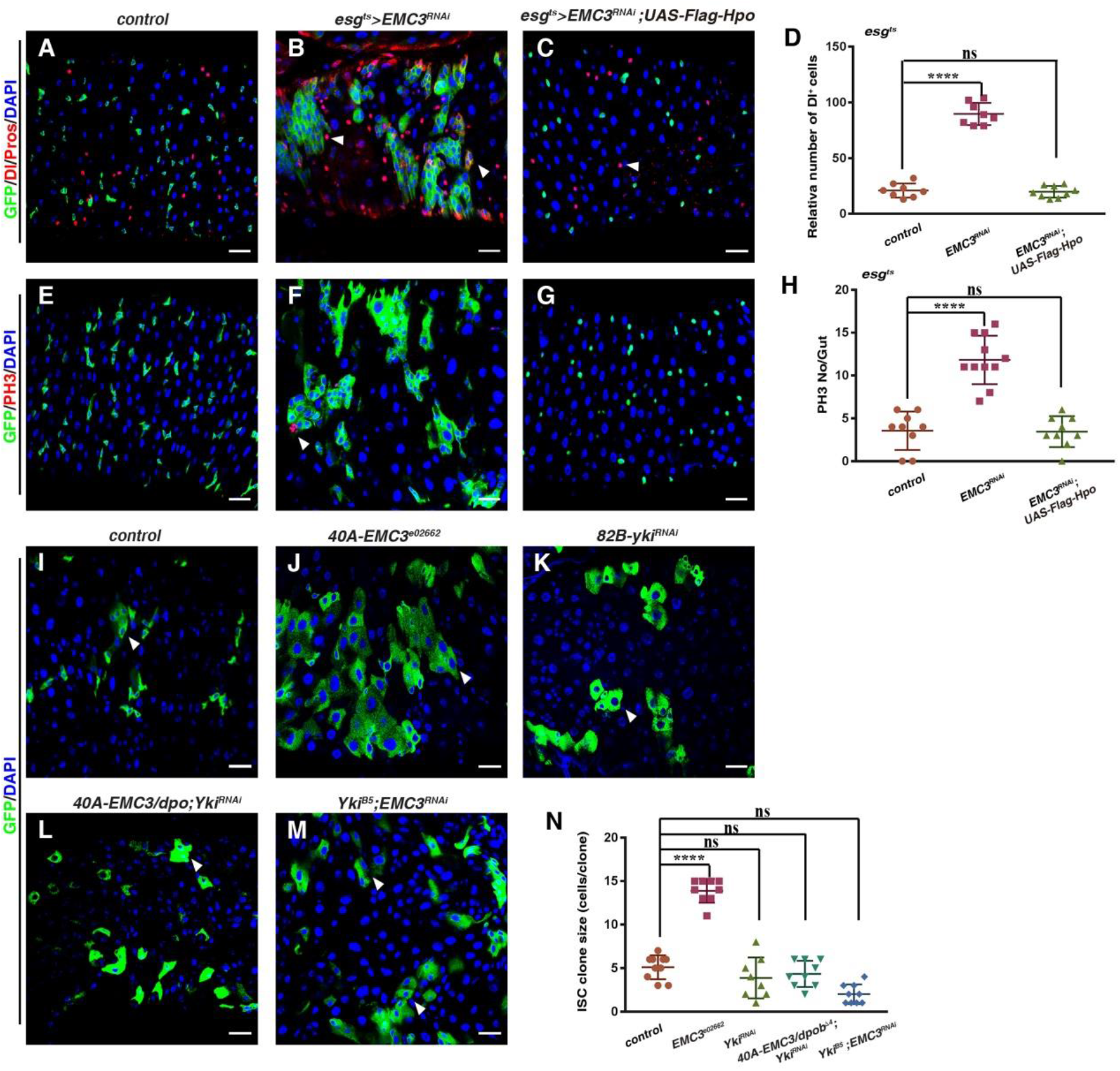
Genetic Interactions between Hpo Signaling and EMC. (A-C) Midguts of the following genotypes stained with DAPI (blue) and anti-Dl/Pros (red): *esg^ts^>w^RNAi^* (A, control), *esg^ts^>EMC3^RNAi^* (B), and *esg^ts^>EMC3^RNAi^; Flag-Hpo* (C). (D) Quantification of the Dl^+^ cell number in intestines with indicated genotypes. Mean ± SD is showed. n ≥ 8. ^ns^*p* > 0.05, *\*\*\*\*p* < 0.0001. (E-G) Midguts of the following genotypes stained with DAPI (blue) and anti-PH3 (red): *esg^ts^>w^RNAi^* (E), *esg^ts^>EMC3^RNAi^* (F), and *esg^ts^>EMC3^RNAi^; Flag-Hpo* (G). (H) Quantification of the PH3^+^ cell number in intestines with indicated genotypes. Mean ± SD is showed. n ≥ 9. ^ns^*p* > 0.05, \*\*\*\**p* < 0.0001. (I-M) MARCM clones co-stained with DAPI (blue): WT clone (I), *EMC3^e02662^* clone (J), *Yki^RNAi^* clone (K), *EMC3/dpob^Δ4^; Yki^RNAi^* clone (L), and *Yki^B5^; EMC3^RNAi^* clone (M). (N) Quantification of cell number per clone in intestines with indicated genotypes. Mean ± SD is showed. n ≥ 8. ^ns^*p* > 0.05, \*\*\*\**p* < 0.0001. Scale bars, 20 μm.

### qRT-PCR Primers Used

*RpL11*-fwd: GGTCCGTTCGTTCGGTATTCGC

*RpL11*-rev: GGATCGTACTTGATGCCCAGATCG

*EMC3*-fwd: AGAAGAAGGCCGAGATCACC

*EMC3*-rev: CGCTGAGATACTTGCCGTTC

*hpo*-fwd: CTCATCGCGCTCAACAATCA

*hpo*-rev: TCAACATAGGCGTGCGATTG

